# Genomic Exploration of Within-Host Microevolution Reveals a Distinctive Molecular Signature of Persistent *Staphylococcus aureus* Bacteraemia

**DOI:** 10.1101/273904

**Authors:** Stefano G. Giulieri, Sarah L. Baines, Romain Guerillot, Torsten Seemann, Anders Gonçalves da Silva, Mark Schultz, Ruth C. Massey, Natasha E. Holmes, Timothy P. Stinear, Benjamin P. Howden

## Abstract

**Background:** Large-scale genomic studies of within-host evolution during *Staphylococcus aureus* bacteraemia (SAB) are needed to understanding bacterial adaptation underlying persistence and thus refining the role of genomics in management of SAB. However, available comparative genomic studies of sequential SAB isolates have tended to focus on selected cases of unusually prolonged bacteraemia, where secondary antimicrobial resistance has developed. To understand the bacterial genomic evolution during SAB more broadly, we applied whole genome sequencing to a large collection of sequential isolates obtained from patients with persistent or relapsing bacteraemia.

**Results:** We show that, while adapation pathways are heterogenous and episode-specific, isolates from persistent bacteraemia have a distinctive molecular signature, characterised by a low mutation frequency and high proportion of non-silent mutations. By performing an extensive analysis of structural genomic variants in addition to point mutations, we found that these often overlooked genetic events are commonly acquired during SAB. We discovered that IS*256* insertion may represent the most effective driver of within-host microevolution in selected lineages, with up to three new insertion events per isolate even in the absence of other mutations. Genetic mechanisms resulting in significant phenotypic changes, such as increases in vancomycin resistance, development of small colony phenotypes, and decreases in cytotoxicity, included mutations in key genes (*rpoB, stp, agrA*) and an *IS256* insertion upstream of the *walKR* operon.

**Conclusions:** This study provides for the first time a large-scale analysis of within-host evolution during invasive *S. aureus* infection and describes specific patterns of adaptation that will be informative for both understanding *S. aureus* pathoadaptation and utilising genomics for management of complicated *S. aureus* infections.

## BACKGROUND

The outcome of *Staphylococcus aureus* bacteraemia (SAB) is a result of a complex interaction of host, pathogen and treatment factors. Persistence, usually defined as bacteraemia of greater than 3-7 days duration, is an important factor in SAB outcome [1], including secondary antibiotic resistance development, metastatic infectious complications, and mortality [2]. Persistent bacteraemia involves a sequence of events, including invasion, immune evasion, and establishment of secondary infectious foci, usually all in the context of antimicrobial treatment [3]. From the bacterial perspective, invasive *S. aureus* isolates are subjected to the pressures of the immune response, lack of nutrients, and antibiotics. These environmental challenges constitute a significant selective pressure driving adaptive evolution in the pathogen, and access to sequential isolates from patients with persistent SAB offers the opportunity to understand pathoadaptation during invasive *S. aureus* infections.

Over the last decade, with increasing availability of whole-genome sequencing, within-host genomic studies have addressed *S. aureus* niches that are important for pathogenesis [4]. Studies of colonising isolates have uncovered the “cloud of diversity” of *S. aureus* colonising the host and improved our ability to track transmission networks [5]. Other authors have revealed mutations associated with the evolution from colonising to invasive strain in a single patient [6] or in large cohorts [7]. Genomic studies of sequential blood isolates in persistent SAB have primarily focused on cases with significant phenotypic changes that might arise in persistent infection, such as secondary resistance to antibiotics (especially the vancomycin intermediate phenotype [8, 9], daptomycin resistance [10]), development of the small-colony phenotype [11, 12] or genetic changes associated with extreme cases of persistence [13]. However, these analyses were restricted to a small number of selected cases and thus offer only limited insights on the general pattern of *S. aureus* evolution during SAB. Understanding the typical pattern of within-host evolution during SAB through large-scale investigation of paired isolates will potentially identify shared genomic signatures associated with *S. aureus* adaptation *in vivo* and inform the use of whole-genome sequencing in the management of SAB more broadly. For example, genomic monitoring of SAB could be used to distinguish true relapses from reinfection with a closely related strain, or track mutations associated with persistence or resistance early in the course of the disease, an approach that has been recently demonstrated for lung cancer [14].

To explore genetic changes associated with persistent or relapsing SAB and compare them to those occurring between colonising and invasive isolates, we applied bacterial whole-genome sequencing to a large cohort of SAB, regardless of phenotypic changes. In addition to the commonly investigated mutational variants, we performed a detailed analysis of chromosome structural variants (e.g. large deletions and insertions, insertions of mobile genetic elements) within same-patient strains. This explorative approach uncovered a diverse mutational landscape and a molecular signature distinctive of persistent bacteraemia. Furthermore, we demonstrate for the first time that structural variation represents an important mechanism promoting genetic plasticity within the host, even in the absence of point mutations and insertions and deletions.

## RESULTS

### Population structure of paired isolates from *S. aureus* bacteraemia reveals a broad genetic background

A collection of 130 *S. aureus* isolates from 57 patients was assembled from two multicenter cohorts of SAB (figure 1, panel A and additional file 4: figure S1). We included 50 SAB episodes with at least two blood isolates collected at a minimum of three days apart (50 index isolates, 61 paired invasive isolates). In addition, 12 colonising isolates were collected from 4 episodes already included in the invasive group and 7 supplementary episodes. The median sample delay between index isolate and paired invasive isolate was 8 days (interquartile range [IQR] 5-23). The clinical context of the paired invasive isolate was persistent bacteraemia (n = 31), relapse on treatment (n = 10); and relapse after treatment (n = 10). Among colonising strains, seven were collected before or at the same time as the index sample (median sampling delay 13 days before index, IQR 1.5-71.5) and five were collected afterwards (median delay 5 days, IQR 1-8).

**Figure 1.**
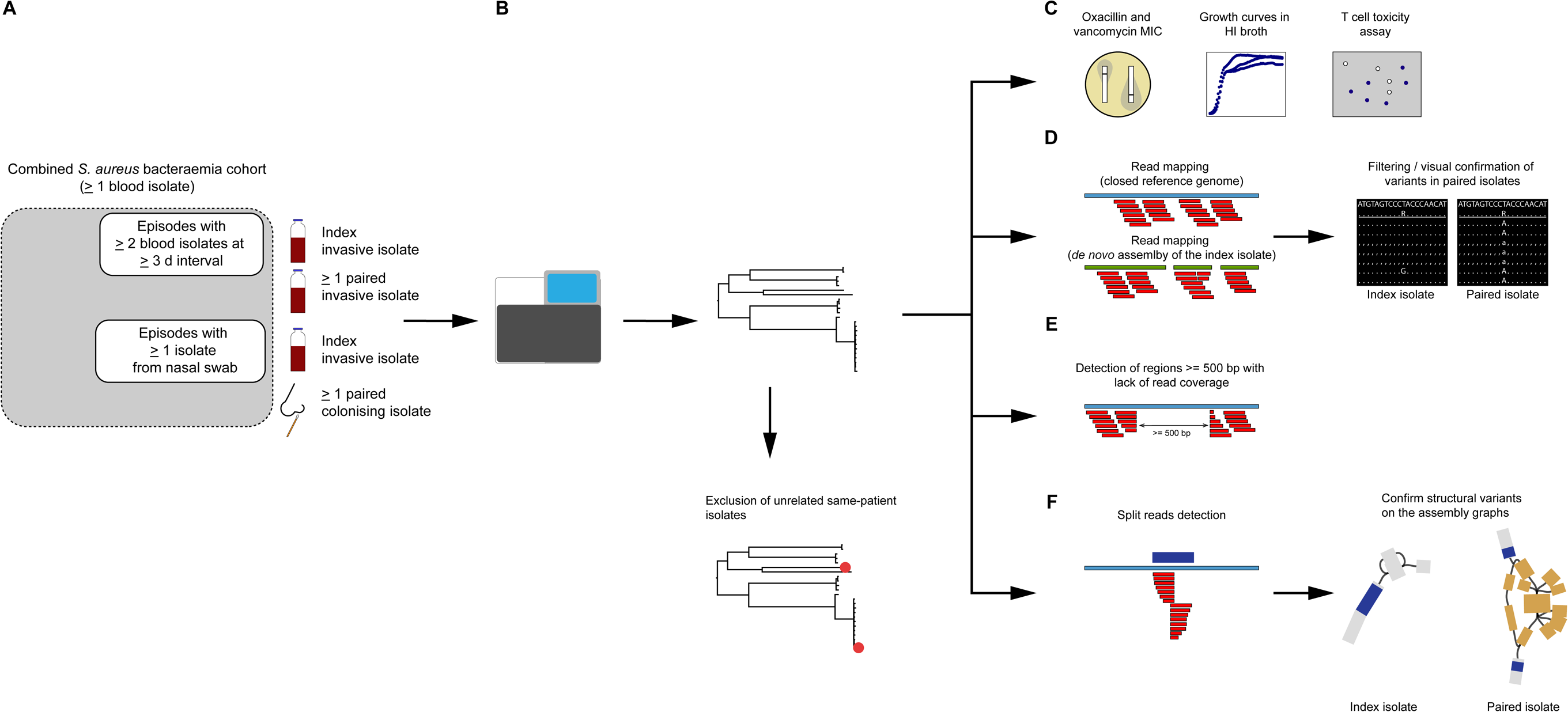
Overview of the study methods. (A) Episodes with at least two blood isolates at least 3 days’ apart, and episodes with at least one isolate from a nasal swab were selected from a combined cohort of *S. aureus* bacteraemia cohort. (B) DNA was extracted from one single colony. Reads from whole genome sequencing were mapped to the reference genome *S. aureus* TW20. Unrelated same-patient isolates (based on SNP distance) were excluded from further analysis. (C-F) Episode-specific phenotypic and genomic analysis. Phenotypic tests included oxacillin and vancomycin MIC, measured by E-test, and overnight growth curves in HI broth (C). Variants calling for SNPs and short indels was performed by mapping on the closes available complete genome and the *de novo* assembly of the index isolate. Variants were filtered based on read depth (≥10) and fraction of reference alleles (>0.5) in the index isolate reads and confirmed by manual inspection of the alignments (D). To identify regions of genome loss that were unique within episode isolates, we scanned the read alignment to the complete genome for intervals with at least 500 bp read coverage loss (E). Screening for structural variants was performed by detecting split reads (along the alignment to the complete genome) that were unique within episode isolates. Structural variants were annotated and confirmed by blasting split intervals on the assembly graph of the episode isolates.

The maximum-likelihood phylogeny of the collection was inferred from 103,974 core genome SNPs and is shown in figure 2A. The population was dominated by CC8, which represented 36% of isolates, followed by CC45 (14%), CC5 (13%), CC22 (8%) and CC39 (7%). The dominant clade was ST239, including four closely related strains with novel ST types that are single-locus variants of ST239. This clade accounted for 28% of the isolates. This diverse population of *S. aureus* shows that the paired isolates were selected from a broad genetic background.

**Figure 2.**
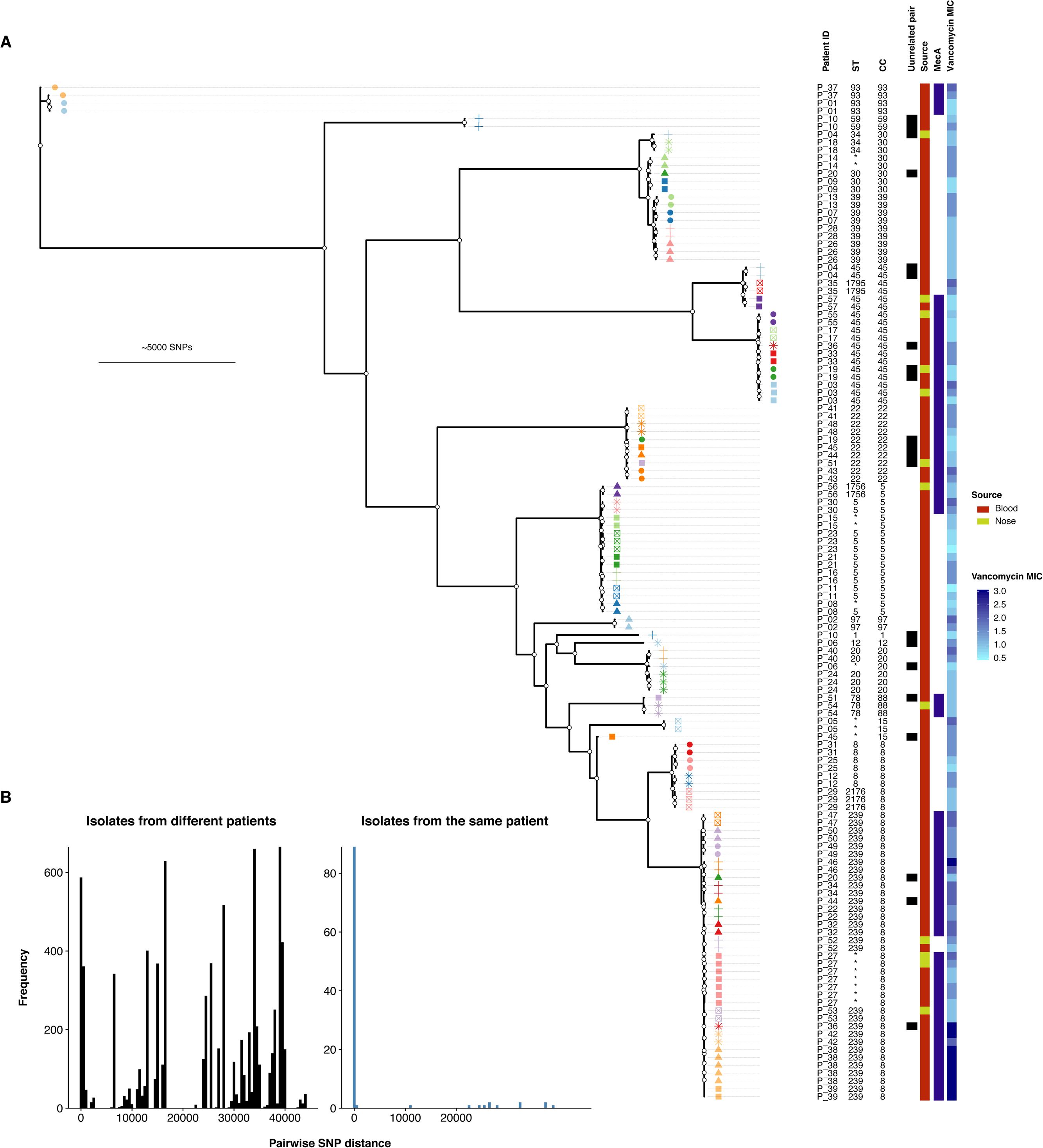
(A) Maximum-likelihood tree of 130 isolates from 57 patients with *Staphylococcus aureus* bacteremia. Patient-specific shape-color combinations annotate branch tips. White circles indicate nodes with ≥ 95% ultrafast support and ≥ 80% SH-like approximate likelihood ratio test support. Frequency distribution of pairwise single-nucleotide polymorphism (SNP) distance between all isolates (B) and isolates from the same patient (C).

We then calculated the pairwise SNP distance and used phylogenetic clustering to infer relatedness of paired isolates and thus distinguish between persistent or relapsing bacteraemia and co-infection or reinfection with an unrelated strain (figure 1, panel B). We also investigated the genetic distance between index blood isolates and their paired colonising isolate to identify SAB episodes that were unrelated to the sampled colonising isolate. Most same-patient isolates clustered together and exhibited a pairwise SNP distance below 100 (figure 2, panel B and C). In this group, pairwise distances ranged between 0 and 98 SNPs, except for one pair, that was separated by 717 SNPs. This latter pair was categorised as related despite the large pairwise distance, because the two ST93 isolates clustered together on the tree. Nine paired isolates (seven paired invasive and two paired colonising) had a SNP distance to the index larger than 1000 bp, and were also different by multilocus sequence type (MLST). We therefore defined these isolates as genetically unrelated to the index and excluded them from further analysis of *in vivo* evolution. The seven unrelated paired invasive isolates were collected after a longer interval as compared to isolates that were genetically close to the index sample (median sampling delay 72 vs. 7 days, p = 0.002, figure 3). Thus, reinfection with a different clone as defined by genetic unrelatedness occurred in 7 out of 50 (14%) cases of SAB included in this study. This is consistent with a previous publication by Fowler *et al.*, where 20% of SAB recurrences were reinfections, as defined by pulsed-filed gel electrophoresis (PFGE), a technique that has lower resolution than WGS [15].

**Figure 3.**
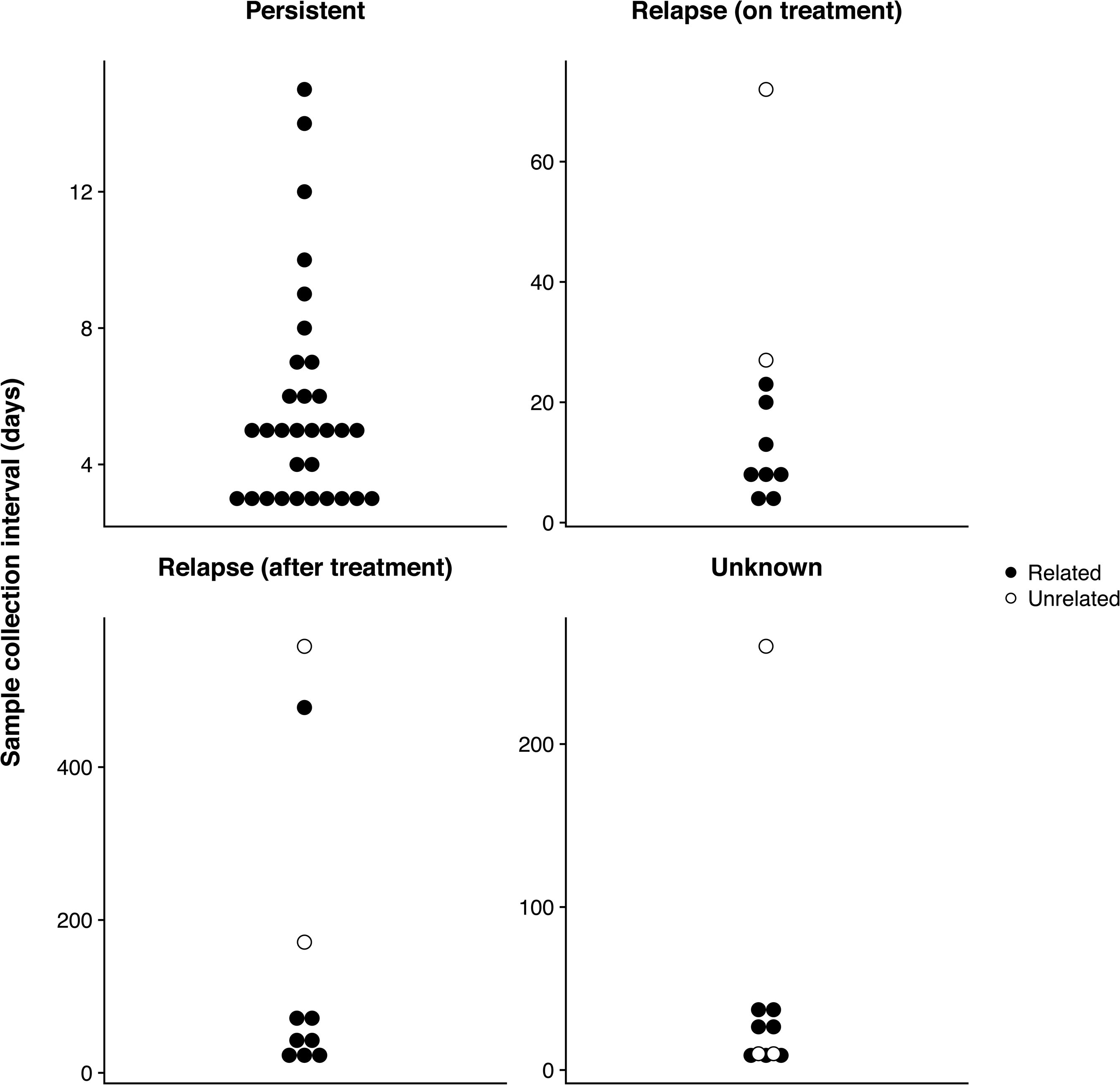
Sample collection interval between index isolate and paired invasive isolate according to the clinical context of the paired isolate.

After exclusion of unrelated isolates pairs, 51 episodes with 115 isolates were retained for in-depth genomic and phenotypic within-host evolution analysis (figure 1, panels C-F).

### Paired invasive isolates have low genetic diversity

The choice of the reference genome and the filtering of variants has an impact on the number of identified mutations [16]. Therefore, to obtain the most accurate estimate of within host diversity in paired isolates, we applied an episode-specific genome mapping approach. By mapping sequence reads to both the closest available complete genome from the NCBI repository and a *de novo* polished assembly of the index isolate and thorough review of the variants through manual inspection of the alignement, we were able to effectively eliminate a significant number of false-positive mutation calls and retain only true genetic variation (additional files 5 and 6:figures S2 and S3). Using this approach, we identified a total of 182 variants (141 SNPs and 41 indels) in 32 out of 64 paired isolates. We observed very limited genetic diversity in paired invasive isolates compared to paired colonising isolates (figure 4, panel A and B). Only 23 (43%) of 54 paired invasive isolates exhibited at least one mutation, while nine out of ten paired colonising isolates were mutated (p = 0.016). Among isolates with at least one mutation, the median number of variants in paired invasive and paired colonising isolates was 2 and 12, respectively (p = 0.014).

**Figure 4.**
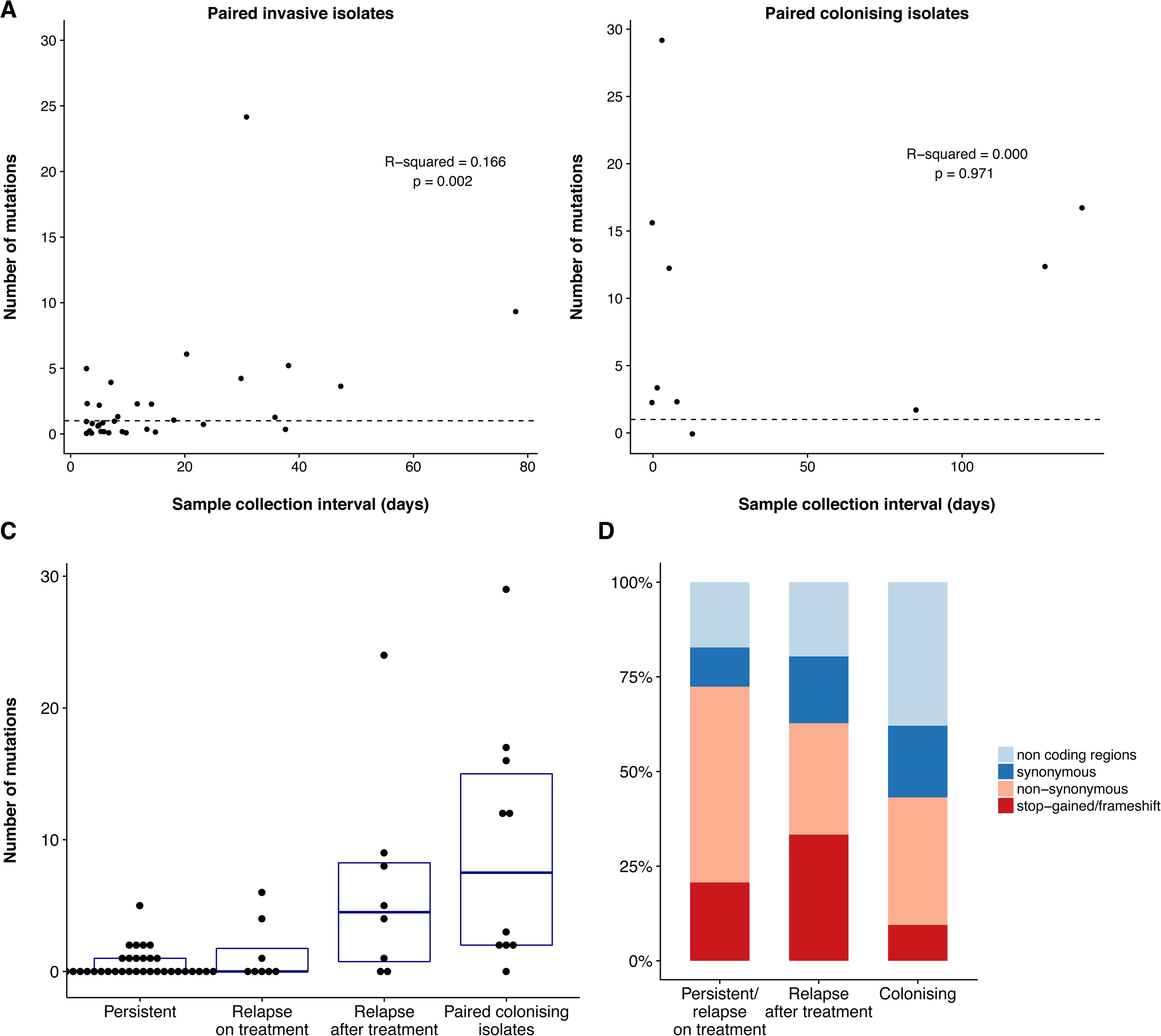
Variants identified by episode-specific mapping and variant calling (after exclusion of unrelated same-patient isolates). Correlation between sample collection interval and number of mutations separating the paired isolates from the index isolate for invasive paired isolates (A) and paired colonising isolates (B), after exclusion of one outlier (isolate collected 478 days after the index). The dotted line represents one mutation. (C) Number of mutations according to the clinical context of paired isolate (persistent bacteraemia, relapse on treatment or relapse bacteraemia after treatment, paired colonising isolate). (D) Distribution of mutation types according to the clinical context of the paired isolate.

Among 158 unique variants, 81 (51%) were predicted to result in changes in protein function: 60 were missense substitutions, 5 were nonsense substitutions (leading to a premature stop codon), and 16 were frameshift mutations. The remaining 77 mutations occurred in non-coding regions (47), or were synonymous substitutions (30) (additional file 2: table S2).

### Colonising isolates and late relapses have a distinctive molecular signature

While the rate of mutation in *S. aureus* may be dependent on the genetic background [17], it is unknown whether evolution rates are different during invasive infection, where host immune response and antibiotic treatment exert a strong selective pressure. We therefore explored associations between mutation counts and clinical, phenotypic and genetic characteristics of the paired isolates. No association was found between mutation count and MRSA status or clonal complex. Interestingly, while there was a weak correlation between length of the collection interval of invasive isolates and mutation count (figure 4A), the association was not linear, with an increase in mutation counts when the collection interval exceeded 15 days (additional file 7: figure S4). Since 15 days is the usual duration of treatment of uncomplicated SAB [18], this suggests that genetic diversity was higher when the paired isolate was collected after treatment. Consistent with this observation, we found a significantly higher number of mutations in paired invasive isolates from relapses after completion of anti-staphylococcal treatment (median 4.5 mutations per isolate) as compared to isolates from persistent bacteraemia or relapses on treatment (median 0 mutations) (Figure 4, panel C). In terms of genetic diversity, isolates from relapses after treatment were as genetically diverse as paired colonising isolates compared to index isolates (figure 4, panel C), indicating that they might represent reinfection with a closely related strain (in other words a new invasive event from the colonising compartment) rather than the result of a persistent invasive focus.

A similar pattern was discovered when we analysed the predicted mutation effects on the encoded proteins. The proportion of non-silent mutations (either nonsynonymous or stop-gained or frameshift) decreased progressively from 72% in invasive isolates from persistent bacteraemia or relapse on treatment to 63% in isolates from relapse after treatment to 43% in colonising isolates (figure 4, panel D). The high proportion of non-silent mutations (66% vs. 43%, p = 0.002) indicates that the invasive compartment may be under stronger positive selection compared to the “colonising compartment”. On the other hand, mutations found in late relapses might arise in the colonising compartment rather than during invasive infection.

### Adaptation pathways are episode-specific

To identify possible convergence of mutation pathways among the 82 variants associated with predicted change in protein sequences, we applied protein sequences clustering using CD-HIT (additional file 3: table S3). Overall, mutation pathways were highly diverse and episode-specific. The only protein-coding gene that was mutated in more than one episode was the accessory gene regulator component *agrA*, with a nonsynonymous SNP in a paired invasive isolate (T88M) and a frameshift in a colonising isolate (at position 127). Given the weak convergence among mutated genes, we attempted to identify common pathways of within-host microevolution by categorising the mutated proteins using the Clusters of Orhologous Groups (COG) database and performing an enrichment analysis using reference genome *S. aureus* TW20 as a comparator. Analysis of 17 categories didn’t show any significant enrichment. Nevertheless, genes related to cell wall and membrane biogenesis among paired invasive isolates reached the lowest p value (uncorrected p value 0.063, additional file 8: figure S5). Overall, different pathways were affected by mutations in invasive and colonising pairs, an observation that is consistent with distinctive selective pressures in the nasal and the blood compartment.

### Mutations in pairs with observed phenotypic changes

#### Antibiotic resistance and growth rate

Within-host phenotypic adaptation might indicate diversifying selection under the selective pressure of antibiotics and the immune system [19]. Therefore, we identified pairs with changes in specific phenotypes between the index isolate and paired isolates. We selected invasive pairs, as they were associated with a stronger positive selection signature with a higher proportion of non-silent mutations. We performed a pairwise analysis of phenotypes that were predicted to change in response to antibiotic pressure (vancomycin MIC and oxacillin MIC, growth rate) (additional file 9: figure S6). Significant changes in phenotypes were observed in a small number of episodes (increase in vancomycin MIC of ≥ 1 µg/ml, 3 episodes, sharp decrease in growth rate leading to small colony variants [SCV], 3 episodes). Details of these episodes are shown in table 1.

**Table 1.** Summary of episodes with phenotypic changes

| Patient code | Type of isolate | Collection interval (days) | Context of paired isolate | Days of treatment | Principal antibiotic | *mecA* | Van MIC (µg/ml) | Median doubling time | Mutations (SNP/indels) | Non-silent mutations |
| --- | --- | --- | --- | --- | --- | --- | --- | --- | --- | --- |
| *Smal-colony variant and vancomycin MIC increase* |  |  |  |  |  |  |  |  |  |  |
| P_03 | Index | - | - | - | - | Pos | 0.75 | 34.5 | - | 0 |
| P_03 | Paired invasive | 38 | Relapse (after treatment) | 14 | Van | Pos | 2 | 53.9 | 4 (1/3) | 4 |
| P_05 | Index | - | - | - | - | Neg | 1.5 | 38.4 | - | 0 |
| P_05 | Paired invasive | 18 | Relapse (after treatment) | 12 | Flx | Neg | 2 | 50.6 | 1 (1/0) | 1 |
| P_30 | Index | - | - | - | - | Pos | 1.5 | 42.1 | - | 0 |
| P_30 | Paired invasive | 14 | Persistent | 14 | Van | Pos | 2 | 83.5 | 2 (1/1) | 2 |
| *Vancomycin MIC increase* |  |  |  |  |  |  |  |  |  |  |
| P_42 | Index | - | - | - | - | Pos | 2 | 44.7 | - | 0 |
| P_42 | Paired invasive | 9 | Persistent | 8 | Van | Pos | 3 | 46.5 | 0 (0/0) | 0 |
| P_46 | Index | - | - | - | - | Pos | 2 | 50.6 | - | 0 |
| P_46 | Paired invasive | 23 | Relapse (on treatment) | 21 | Van | Pos | 3 | 48.9 | 0 (0/0) | 0 |

Van: vancomycin. Flx: flucloxacillin. MIC: minimum inhibitory concentration. SNP: single nucleotide polymorphism

**Table 2.**
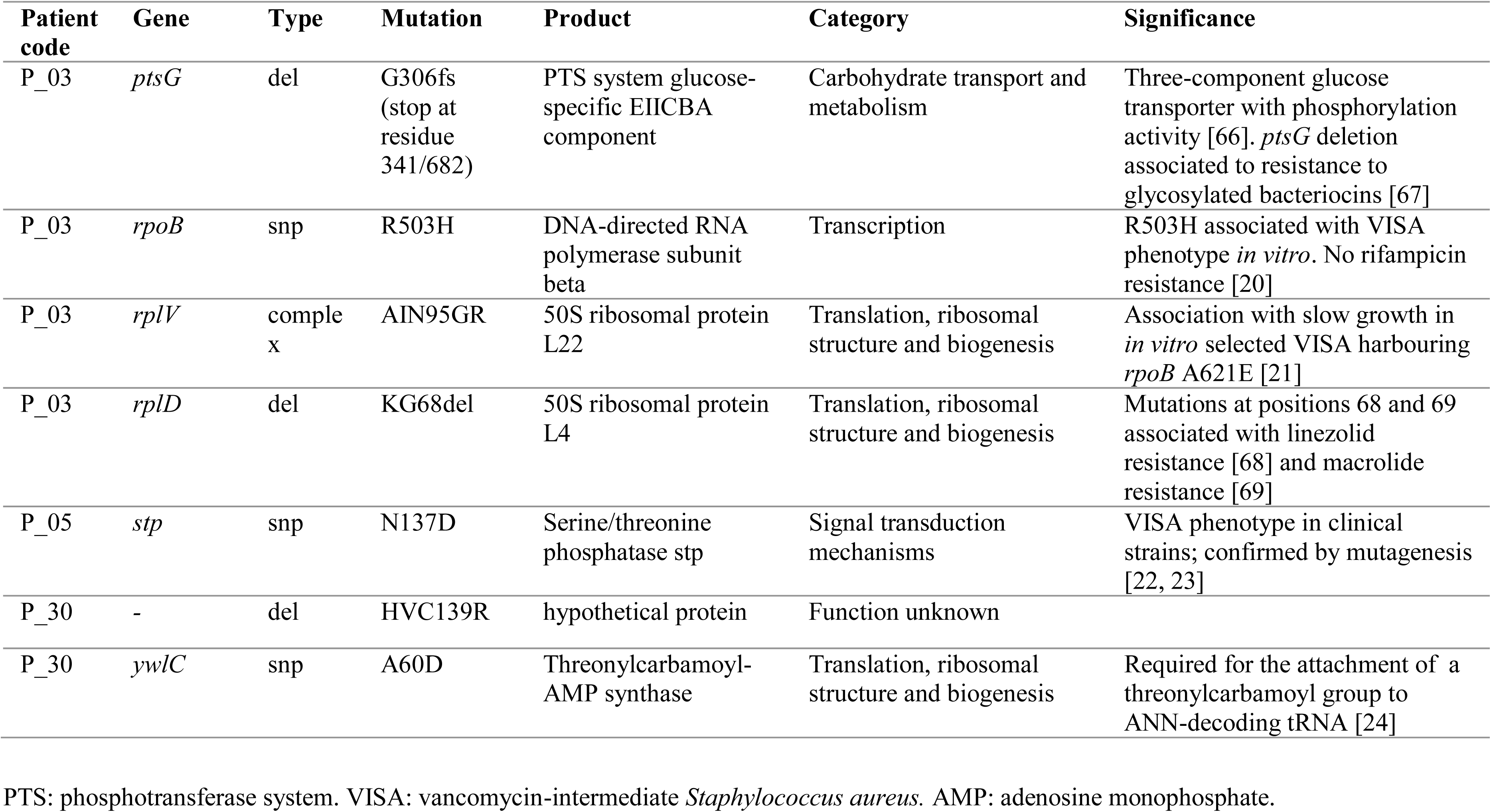
Mutations in episodes with phenotypic changes

| Patient code | Gene | Type | Mutation | Product | Category | Significance |
| --- | --- | --- | --- | --- | --- | --- |
| P_03 | *ptsG* | del | G306fs (stop at residue 341/682) | PTS system glucose-specific EIICBA component | Carbohydrate transport and metabolism | Three-component glucose transporter with phosphorylation activity [66]. *ptsG* deletion associated to resistance to glycosylated bacteriocins [67] |
| P_03 | *rpoB* | snp | R503H | DNA-directed RNA polymerase subunit beta | Transcription | R503H associated with VISA phenotype *in vitro*. No rifampicin resistance [20] |
| P_03 | *rplV* | complex | AIN95GR | 50S ribosomal protein L22 | Translation, ribosomal structure and biogenesis | Association with slow growth in *in vitro* selected VISA harbouring *rpoB* A621E [21] |
| P_03 | *rplD* | del | KG68del | 50S ribosomal protein L4 | Translation, ribosomal structure and biogenesis | Mutations at positions 68 and 69 associated with linezolid resistance [68] and macrolide resistance [69] |
| P_05 | *stp* | snp | N137D | Serine/threonine phosphatase stp | Signal transduction mechanisms | VISA phenotype in clinical strains; confirmed by mutagenesis [22, 23] |
| P_30 | - | del | HVC139R | hypothetical protein | Function unknown |  |
| P_30 | *ywlC* | snp | A60D | Threonylcarbamoyl-AMP synthase | Translation, ribosomal structure and biogenesis | Required for the attachment of a threonylcarbamoyl group to ANN-decoding tRNA [24] |

PTS: phosphotransferase system. VISA: vancomycin-intermediate *Staphylococcus aureus*. AMP: adenosine monophosphate.

Genetic changes underlying the SCV phenotype and secondary increase in vancomycin MIC were diverse, indicating that phenotypic convergence was not associated with genetic convergence. Patient 3 presented with a ST45-MRSA bacteraemia, which was associated with a dialysis-catheter device and was treated with vancomycin for 14 days. She had recurrent bacteraemia 38 days after the first episode. The recurrent strain exhibited an increase in vancomycin MIC from 0.75 to 2 µg/ml and a SCV phenotype. We identified four mutations arising in the relapsing strain: a non-synonymous SNP in the *rpoB* gene leading to an arginine-histidin substitution in position 503 (R503H), a deletion in position 283 of the *rplV* gene (ribosomal protein 22), a deletion in position 66 of the *rplD* gene (ribosomal protein 4) and a deletion leading to a truncation of gene *ptsG*. The *rpoB* R503H and ribosomal protein mutations have been previously described in *in vitro* selected vancomycin-intermediate mutants [20, 21], but never in clinical isolates. The *rpoB* R503H mutation is not associated with rifampicin resistance, consistent with the lack of exposure to rifampicin in this case. Patient 5 was treated with vancomycin for 3 days and flucloxacillin for 13 days for a ST15-MSSA catheter-related bacteraemia and experienced relapse at day 18. The relapsing strain had a SCV phenotype and vancomycin MIC increased from 1.5 to 2 µg/ml. It had a nonsynonymous SNP in the serine/threonine phosphatase (*stp*) gene leading to a N137D mutation. *stp* mutations have been identified previously in persistent SAB with secondary development of VISA [22, 23]. Finally, patient 30 had relapsing ST5-MRSA bacteraemia after 14 days of vancomycin (combined with rifampicin and ciprofloxacin). Median doubling time of the relapsing strain increased from 42 to 83 min, while vancomycin MIC increased from 1.5 to 2 ug/ml. Mutation analysis identified a deletion in a protein whose function could not be predicted and a non-synonymous SNP (mutation A60D) in the *ywlC* gene, which encodes a translational factor (threonylcarbamoyl-AMP synthase) [24]. This gene has never been linked to vancomycin resistance or growth rate. However, a *ywlC* ortholog has been shown to be essential in *E. coli* [24], thus it is possible that point mutations impair *S. aureus* growth.

In two episodes with an increase in vancomycin MIC by at least 1 µg/ml, no mutation separated the index isolate from the paired isolate, suggesting that other genetic changes may have occurred (see below).

#### Cytotoxicity

Recently, it has been shown that within-host evolution from colonising to invasive *S. aureus* can be associated with a dramatic decrease in cytotoxicity [25], however it is unknown whether a similar trend can be observed during persistent infection. To assess evolution of cytotoxicity during SAB, we tested a subset of 21 episodes that were considered more likely to be associated with changes based on phenotypic characteristics (i.e. longer duration of bacteraemia, relapse on anti-staphylococcal treatment, small colony phenotype or secondary increase in vancomycin MIC) or because of mutations in the *agrA* gene.

Similar to the other pairwise phenotypic tests, cytotoxicity remained unchanged between index isolate and paired isolate, with the dramatic exception of one paired invasive isolate with *agrA* mutation T88M, which was associated with a marked reduction in THP-1 cell lysis (from 56% non-viable cells to 11%) as compared to the index isolate (additional file 5 figure S2).

### Chromosome structural variants

The potential significance of chromosome structural variants in *S. aureus* resistance and adaptation has been recently highlighted [13, 26]. However, when using only partially assembled genomes or read-mapping the characterisation of structural variants is much more challenging than SNP-calling and these types of changes are often overlooked. Using an approach combining read coverage arithmetic, read filtering and annotation of split reads (figure 1), we detected 21 unique structural variants within 15 SAB episodes: two plasmid losses, five large deletions (ranging from 261 to 15,622 bp), one recombination and 13 insertions (summarised in table 3). Beside the two instances of plasmid loss, two large structural changes were particularly interesting. A recombination of prophage Sa phi3 encoding the immune escape cluster (IEC) was identified when comparing the index blood isolate of patient 19 with its paired colonising isolate. As a consequence, the invasive isolate carried the staphylokinase gene (*sak*) and the staphylococcal complement inhibitor gene (*scn*), while the colonising isolate carried the complete IEC including *sak, scn* and the chemotaxis inhibitory protein (*chp*). Since there were no enterotoxin genes, this constellation can be classified as IEC type E and B, respectively, according to the classification proposed by van Wamel et al [27]. In patient 21 we observed a deletion of pathogenicity island SaPi2 in the paired invasive isolate that was collected upon relapse of bacteraemia 65 days after the first episode.

**Table 3.** Overview of chromosome structural variants

| Patient code | Type of variant | Reference (position) | Annotation | Isolate with variant |
| --- | --- | --- | --- | --- |
| P_27 | plasmid loss | *S. aureus* TW20 | 29,585 bp-plasmid containing the chlorhexidine tolerance determinant qacA and cadmium resistance genes | Index, paired invasive, one paired colonising (second paired colonising like reference) |
| P_30 | plasmid loss | *S. aureus* SA564 | 27,272 bp-plasmid containing the beta-lactamase operon | Index (paired invasive like reference) |
| P_05 | deletion | *S. aureus* NCTC_8325 (2188042) | 861 bp-deletion (fructose bisphosphate aldolase) | Paired invasive |
| P_12 | deletion | *S. aureus* NCTC_8325 (1502725) | 261 bp-deletion (phage protein) | Index (paired invasive like reference) |
| P_14 | deletion | *S. aureus* FORC_001 (1953000) | 597 bp-deletion (hypothetical protein) | Paired invasive |
| P_21 | deletion | *S. aureus* SA564 (798999) | 15622 bp deletion of pathogenicity island SaPi2 | Paired invasive |
| P_54 | deletion | *S. aureus* AUS0325 (1191813) | 495 bp-deletion (serine/threonine-protein phosphatase) | Index (paired colonising like reference) |
| P_19 | recombination | *S. aureus* CA-347 (1577530) | Phage Sa phi3 recombination | Index (paired colonising like reference) |
| P_22 | insertion | *S. aureus* TW20 (2243085) | IS256 insertion (prophage phiSPbeta-like) | Paired invasive |
| P_27 | insertion | *S. aureus* TW20 (173671) | IS256 insertion (putative DNA-binding protein) | Index and one paired invasive |
| P_27 | insertion | *S. aureus* TW20 (1223135) | IS256 (putative membrane protein) | Index, paired invasive (paired colonising like reference) |
| P_27 | insertion | *S. aureus* TW20 (1971406) | IS256 insertion (lantibiotic immunity protein, genomic island niSa beta) | Index, paired invasive, one paired colonising (second paired colonising like reference) |
| P_27 | insertion | *S. aureus* TW20 (173671, 173626) | IS256 insertion (putative DNA-binding protein) | Index and one paired invasive |
| P_34 | insertion | *S. aureus* TW20 (1619645) | IS256 insertion (hypothetical protein) | Index (paired invasive like reference) |
| P_39 | insertion | *S. aureus* TW20 (1955676) | IS256 insertion (genomic island niSa beta) | Index (paired invasive like reference) |
| P_46 | insertion | *S. aureus* TW20 (24644) | IS256 insertion (upstream of walKR operon) | Paired invasive |
| P_46 | insertion | *S. aureus* TW20 (528848) | IS256 insertion (exotoxin, genomic island niSa alpha) | Paired invasive |
| P_46 | insertion | *S. aureus* TW20 (2306594) | IS256 insertion (phage protein) | Paired invasive |
| P_47 | insertion | *S. aureus* str._JKD6008 (1914213) | IS256 insertion (genomic island niSa beta) | Paired invasive |
| P_47 | insertion | *S. aureus* str._JKD6008 (2227509) | IS256 insertion (P-ATPase superfamily P-type ATPase potassium (K<sup>+</sup>) transporter subunit A) | Paired invasive |
| P_49 | insertion | S. aureus<br>str._JKD6008<br>(702665) | IS256 insertion (upstream of sodium/hydrogen<br>exchanger family protein) | Paired invasive |
| P_52 | insertion | S. aureus<br>TW20<br>(1955675) | IS256 insertion (genomic island niSa beta) | Paired colonising |

Bp : base pair

The most prevalent structural change in our cohort was the insertion of IS*256* elements. We observed 13 unique IS*256* insertions in 8 episodes. Interestingly, the number of insertions didn’t correlate with the number of mutations, and up to three new IS*256* insertion were found in paired invasive isolates with no mutations relative to index (additional file 10: figure S7).

Intriguingly, all strains with new insertions belonged to ST239 or to a closely-related single-locus variant of ST239. A BLAST search of the 1324bp-long IS*256* sequence among all available *S. aureus* complete genomes and the draft assemblies of the 130 isolates included in our study confirmed that IS256 is highly disseminated in ST239 and restricted to a few other sequence types (additional file 11: figure S8).

We mapped split reads that were unique within single episodes on a single ST239 reference genome (*S. aureus* TW20) and found that there were hotspots for these new *IS256* insertions on the chromosome (figure 5). One of these hotspots was the genomic island niSa beta with unique new insertions in four different patients, two of them around the lantibiotics operon. Moreover, we discovered that the paired invasive isolate from patient 46 had three new IS*256* insertions including one 150 bp upstream of the *walKR* operon. This finding was relevant because the isolate showed an increase in vancomycin MIC from 2 to 3 ug/ml as compared to the index isolate but no point mutations were found (see above). Notably, IS*256* insertion and tempering of WalKR activity has been previously shown to cause VISA phenotype *in vitro*, but has never been described during human infection [26].

**Figure 5.**
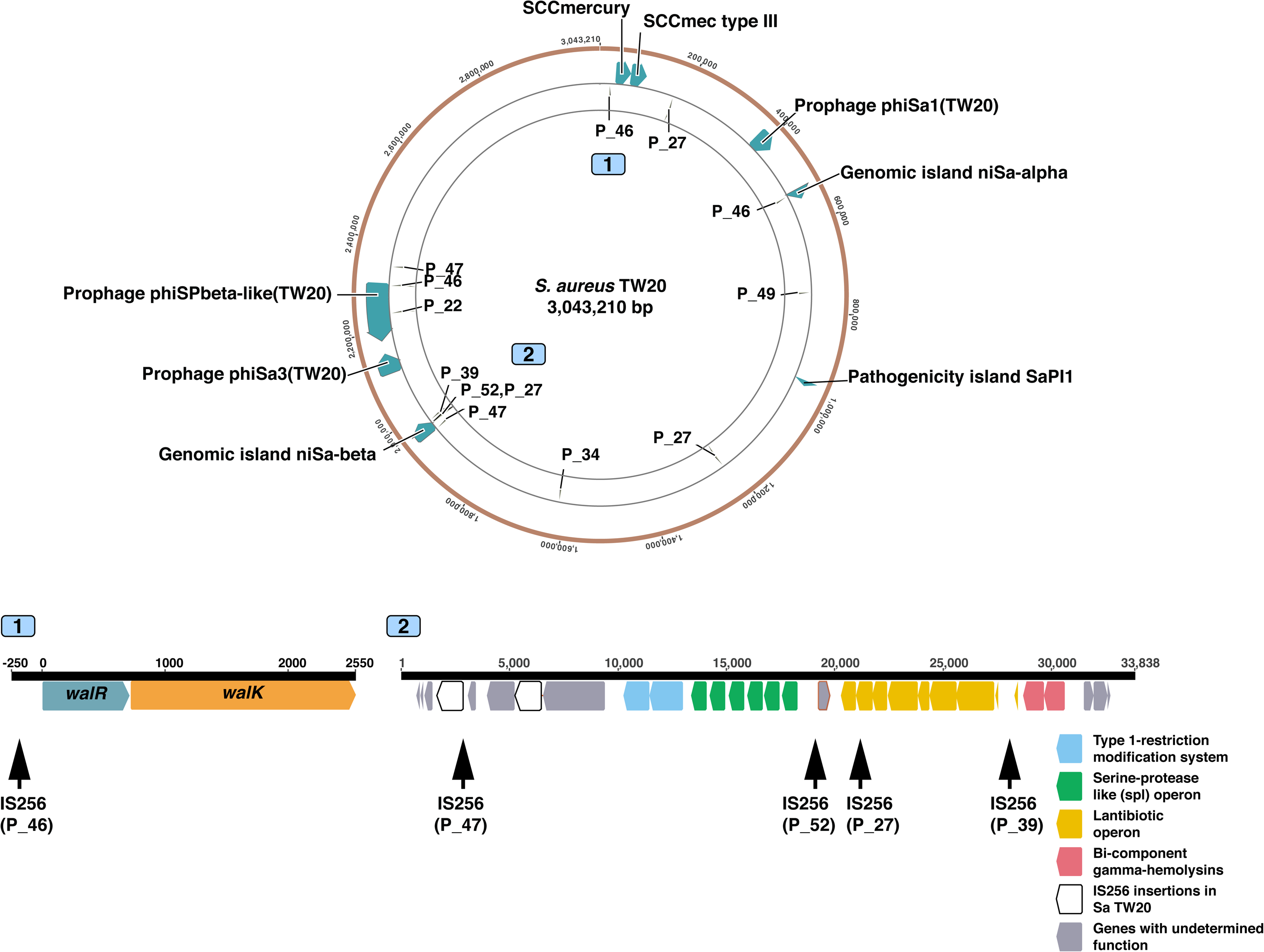
Location of *IS256* insertions differentiating CC239 isolates within the same patient. Insertions are mapped on the chromosome of the reference *S. aureus* TW20 and labelled with the patient ID. Two sites of interest are depicted in detail. The diagram of site 1 (position 24,645-27,443) shows an *IS256* insertion 150 bp upstream of the two-component regulator *walKR,* that was found in the paired invasive isolate but not in the index isolate of patient 46. Site 2 (position 1,950,453-1,984,290) is the genomic island niSa-beta, which appears to be a hotspot of new *IS256* insertions within same-patient isolates.

## DISCUSSION

This large-scale comparative genomics study of patients with persistent or recurrent SAB provides the first comprehensive overview of within-host *S. aureus* diversity associated with bacteraemia. By compiling a curated inventory of mutational and structural within-host variants across different genetic backgrounds and manifestations of SAB, we show that invasive isolate pairs have a specific molecular signature (denoted by limited diversity and a high proportion of non-silent mutations) and that structural variation and especially insertion of IS*256* elements enhances genetic diversity during human infection. With the notable exception of *agrA*, which was mutated in one invasive pair and one invasive-colonising pair, there was no convergence at the gene level among mutations and indels, indicating that pathways of adaption are episode-specific, even when we found common phenotypic changes within pairs. Loss of *agr* function within the host has been previously described both among colonising isolates [5] and during invasive infection [8, 28]. Enrichment for *agrA* mutations was also observed in a study of 105 colonising-invasive pairs [7]. While one of the two mutations was associated with the only significant reduction in cytotoxicity observed in our cohort, none led to an increase in vancomycin MIC, despite the known link between *agr* dysfunction and vancomycin resistance [29].

When considering bacterial within-host diversity in invasive *S. aureus* infections, it is important to keep a “pathogen-centric” perspective and consider a model consisting of two “compartments”, i.e. the colonising compartment (anterior nares, or other mucosal areas) and the invasive compartment (blood and tissue/organs of primary or metastatic *S. aureus* infection). Bacteria in the colonising compartment are subjected to evolution pressures (competition of the nasal microbiota, some immune system control and intermittent antibiotic exposure at low concentration) but also to purifying selection, since colonisation sites such as the nose are ther natural ecological niche of *S. aureus* [19, 30]. By contrast, bacterial invading blood and tissue are subjected to a formidable selective pressure, including antibiotics at high concentration, host antimicrobial peptides, immune cells, sequestration of nutrients (e.g. iron). This is supported by convergent evolution analysis at the gene ontology level. In this study, a non-significant enrichment for mutations in genes associated with cell wall and membrane metabolism was found in paired invasive isolates, while enrichment for genes associated with cell wall and adhesion was described by Young *et al* among colonising-invasive pairs [7].

Since the advent of WGS, studies addressing within-host diversity of *S. aureus* bacteraemia have mainly focused on genetic changes associated with secondary development of the VISA phenotype under vancomycin pressure [8, 9, 11, 13, 22, 31], or with secondary development of daptomycin resistance [10]. By selecting SAB episodes with phenotypic changes, this approach helps to distinguish evolution in the blood/tissue compartment from background diversity of the colonising *S. aureus* population. Our study complements this previous work by providing for the first time a wider picture of within-host diversity in SAB in a diverse genetic background. By using phenotypic tests such as vancomycin susceptibility testing and growth curves, we showed that secondary changes were present in a small proportion of cases. However, even in this group with more evident features of positive evolution, we found a heterogeneity of mutations. This observation, together with the wide range of alleles described in the VISA literature [29], highlights the multiplicity of pathways by which *S. aureus* adapts to vancomycin pressure *in vivo*. Mutations identified in episodes without detected phenotypic changes were also very heterogenous, and thus we were not able to detect convergence in our dataset or identify mutational hotspots that were associated with persistence or recurrence. This clearly shows that we should be careful in drawing general conclusions on *S. aureus* pathoadaptation from mutations identified in single clinical cases.

One striking finding was the combination of limited genetic variability and high frequency of non-silent mutations among invasive pairs. This is consistent with the “bottle neck” hypothesis of SAB, as shown in animal models, where only individual clones among the diverse colonising pool become invasive [32]. Despite the lack of identified molecular hotspots of persistence, this signature (low abundance of mutations and low fraction of “silent mutations”) may help distinguish between reinfection with a close related colonising strain from relapse from a persistent infection focus. On the background of increased availability of WGS, within-host diversity data could be used not only to understand the pathogenesis of SAB and antibiotic resistance, but also to inform clinical management of persistence or recurrence.

Mobile genetic elements (MGE) are key drivers of evolution in *S. aureus* [33]. Evolution experiments *in vivo* have shown an intense exchange of MGE in piglets co-colonised with different lineage of *S. aureus* [34] and there is evidence of phage recombination in studies of same-patient colonising isolates [5, 30]. In addition to MGE, we have recently illustrated that a large chromosome duplication mediated vancomycin resistance and immune evasion in a case of extremely protracted SAB [13]. However, we don’t have an overview of structural variation and MGEs movements during clinical invasive *S. aureus* infection. Therefore, we applied an episode-specific strategy to detect structural variation by carefully assessing read coverage and split reads within the pairs to identify unique structural changes that were confirmed by review of the assembly graphs. This mapping-based approach allowed us to reveal large changes occurring even in the absence of point mutations. Some of these modifications may have a relevant impact on the phenotype. For example, we observed the loss of pathogenicity island SaPI2 in a paired invasive isolate and a recombination of phage Sa phi3 (including the IEC) in an invasive-colonising pair.

While deletion and recombination were episode-specific without a discernible pattern, a striking finding was the remarkable high frequency of new IS*256* insertions within the same episode among strains belonging to the dominant lineage ST239 (at least one insertion event in eight out of thirteen episodes) with the genomic island niSa beta as a hotspot of new insertions. This genomic island is enriched with IS*256* that has been shown to engender chromosomal inversion in an ST8-IV MRSA [35]. The effect of the new IS*256* insertions in genomic island niSa beta in our paired clinical isolates are uncertain, although insertions around the lantibiotic operon could be important in modulating the production of lantibiotics depending on whether *S. aureus* is in the colonising compartment (i.e. in competing with other microbiota) or is invasive [36]. Up to three new insertions occurred in paired isolates even in the absence of point mutations, suggesting that IS*256* is an efficient mechanism of genetic variability in the environment of invasive *S. aureus* infection,characterised by high selective pressure and reduced effective population size, as it is known that bacterial stress like antibiotic exposure activates insertion sequences [37, 38]. A paradigmatic example was the insertion of IS*256* upstream of the *walKR* operon (in an isolate whose vancomycin MIC increased from 2 to 3 ug/ml), a mechanism that we and others have previously elucidated in *in vitro* selected VISA strains [26, 39].

Our study has some limitations. Because patients were recruited at detection of positive blood cultures for *S. aureus*, colonising *S. aureus* strains were available only for a very small proportion of patients. Therefore, we included invasive-colonising pairs as a comparator to invasive pairs, but our dataset prevents conclusions on molecular signatures on “invasiveness” of *S. aureus*. Furthermore, in our study one colony per sample was sequenced. Recent work has exposed the diversity of colonising *S. aureus* strains by sequencing multiple colonies per sample (up to 12) [5]. Data obtained from this “high resolution” approach can then be used to better infer within-host phylogenies and shed light into the pathogenesis of SAB. Additionally, we tested only a limited array of phenotypes. Data on more complex phenotypes (e.g. immune evasion) might have furnished additional insights into the impact of the host immunity on *S. aureus* evolution within the blood or tissue compartment. Finally, since our analysis of structural variants is based on short read data, we may have missed structural changes that can be usually only detected by long-read sequencing, such as chromosomal inversion.

## CONCLUSIONS

By applying comparative genomics to 57 episodes of SAB with sequential invasive *S. aureus* isolates or paired colonising isolates, we describe specific patterns of *S. aureus* evolution within the invasive compartment (in particular limited within-host diversity and strong positive selection signatures), demonstrate the multiplicity of adaptive changes under the combined pressure of antibiotics and host immunity and highlight the crucial role of structural changes and in particular MGEs like insertion sequence during microevolution within the host. Data from this study will improve our understanding of bacterial pathogenesis SAB and contribute to defining a molecular signature of persistence / relapse that might be used for both biological research and infection management.

## METHODS

### Case and isolate selection

Isolates included in this study were selected from two multicenter cohorts of SAB (figure 1, panel A). The vancomycin substudy of the Australian and New Zealand Cooperative on Outcome in Staphylococcal Sepsis (ANZCOSS) study was a retrospective study of *S. aureus* isolates collected between 2007 and 2008 [40-44]. The Vancomycin Efficacy in Staphylococcal Sepsis in Australasia (VANESSA) cohort was a prospective, multicentre study that was been designed to establish the impact of host, pathogen and antimicrobial factors on outcome from SAB and has recruited patients between 2012 and 2013 [45]. Both studies collected data on patient demographics, comorbidities, clinical characteristics, anti-staphylococcal treatment, and duration of bacteraemia, 30 day-mortality and SAB recurrence. Isolates from subsequent positive blood cultures and from nasal colonisation screening were available for a subset of SAB episodes. Therefore, SAB episodes with at least two blood isolates collected at a minimum of three days apart were included as *invasive* episodes, and the isolate collected at the detection of bacteraemia was defined as *index* isolate; blood isolates collected subsequently were defined as *paired invasive*. Episode for which colonising *S. aureus* isolates were available were included as *colonising-invasive pairs* (*index* isolate: first detected blood isolate; *paired colonising* isolate). Invasive isolates collected after the index isolate were classified according to the clinical context in: (i) *persistent bacteraemia* (no negative blood cultures before the collection of the paired isolate); (ii) *relapse on treatment* (at least one negative blood culture between the index isolate and the paired isolate and collection before the end of antistaphylococcal treatment of the index episode); (iii) *relapse after treatment* (collection after the end of antistaphylococcal treatment of the index episode).

The first blood culture isolate from each episode (index isolate), isolates from blood cultures collected > 3 days after the index (paired isolates) and colonising isolates were stored at −80°C. Phenotypic confirmation of *S. aureus* was performed using the coagulase and DNase tests.

### Whole genome sequencing

Bacterial isolates stored in glycerol broth at −80°C were subcultured twice onto horse blood agar. Genomic DNA was extracted from single colonies using the Janus^®^ automated workstation (PerkinElmer) or manually using Invitrogen PureLink genomic DNA kit or the Sigma GenElute kit. DNA concentration was measured using the Qubit^®^ dsDNA HS Assay Kit (Life Technologies) and normalized to a concentration of 0.2 ng/µl for library preparation with Nextera^®^ XT DNA (Illumina). Genome sequencing was carried out on the MiSeq^®^ and NextSeq^®^ (Illumina) platforms with a read length of 2 × 150 bp or 2 × 250 bp (figure 1, panel B). The quality of sequencing was evaluated by calculating mean read depth (based on a genome length of 3 million bp), and assessing metrics obtained using Spades, version 3.9.0 [46]. Species was confirmed by k-mer classification using Kraken, version 0.10.5-beta [47].

### Multi-locus sequence typing and resistome

*De novo* assemblies of the isolates were generated with Spades [46]. Assembled genomes were scanned for MLST typing using MLST, version 2.7 (T. Seemann, https://github.com/tseemann/mlst). Resistance genes were detected from assemblies using Abricate, version 0.3 (T. Seemann, https://github.com/tseemann/abricate) using the ResFinder database [48]. Clonal complexes were inferred using eBurst, version 3 [49].

### Global core genome alignment

To obtain a global alignment of all isolates included in the study (both invasive and colonising), sequence reads were mapped to *S. aureus* TW20, a clonal complex (CC) 8/ sequence type (ST) 239 methicillin-resistant *S. aureus* (MRSA) reference genome (figure 1, panel B). [50]. Read mapping, variant calling and core genome alignment were performed using the Snippy pipeline, version 3.0 (T. Seemann, https://github.com/tseemann/snippy). Maximum likelihood phylogeny was obtained using IQ-TREE, version 1.6 [51, 52]. Branch support was calculated using both ultrafast bootstrap support [53] and the SH-like approximate likelihood ratio test [54] with threshold values of 95% and 80%, respectively. The phylogenetic tree was plotted and annotated with the R packages *ape* [55] and *ggtree* [56]. The pairwise SNP distance matrices of isolates from different patients and same-patient isolates were compared to determine relatedness between same-patient isolates. Only related same-patient isolates were kept for further phenotypic and genomic analysis (episode-specific analysis).

### Episode-specific analysis. *Phenotypic testing*

Vancomycin and oxacillin MIC were assessed using Etest (bioMerieux), according to manufacturer’s instructions. For growth curves, isolates freshly subcultured were grown overnight in heart infusion (HI) broth, inoculated into 200 µl of fresh HI at a 1:400 dilution, and incubated at 37°C with agitation during 16 hours. Optical density at 600 nm was measured at 15 min interval using the EnSight^TM^ Multimode Plate Reader (PerkinElmer).

Cytotoxicity assays were performed on a subset of isolates that were selected using the following criteria: bacteraemia duration of at least 7 days, relapse on anti-staphylococcal treatment, vancomycin MIC increase or development of small-colony phenotype, possible change in toxicity based on genetic changes (e.g. *agr* mutations).

Cytotoxicity was measured using a modified method of that described previously [57, 58]. A single bacterial colony was inoculated into 5 mL brain heart infusion (BHI) broth (Oxoid), and incubated for 18 hours at 37° C with agitation (180 rpm). A 1 mL aliquote was centrifuged (10 mins, 10,000 rpm), the supernatant collected and frozen at −20°C. Instead of T2 cells, a THP-1 human moncyte cell line was used. THP-1 cells were cultured in RPMI (Lonza) supplemented with 10% fetal calf serum, and 1% L-Glutamine (200mM) – Penicillin (10,000 units) – Streptomycin (10mg/mL) solution (Sigma) at 37°C for 2 to 4 days. For testing, THP-1 cells were centrifuged (10 mins, 1200 rpm, 22°C), and resuspended in Dulbecco’s Phosphate Buffered Saline (Thermo-Scientific) to a concentration of 2.4 - 3.0 million cells/mL. Bacterial supernatant (diluted 50% in BHI broth) and THP-1 cells were mixed in a 1:1 ratio (using 20ul volumes) and incubated for 12 mins at 37°C. After incubation, 20ul of Trypan Blue Solution (Corning) was added and 20ul was loaded onto a disposable cell counting slide (Immune Systems). Three 4×4 grids were counted and averaged to gain a viable- and total-cell count. Each bacterial supernatant was tested in technical duplicate. In the case of the paired invasive isolates (BPH3706 and BPH3757) where a difference in cytotoxicity was detected, two additional biological replicates (each tested in technical duplicate) were performed.

#### Variant calling

We used Snippy to map sequence reads from the same episode to the closest available complete genome in the NCBI repository (strain names and accession numbers are listed in additional file 1: table S1) and to the *de novo* assembly of the index isolate, which was generated using Spades as described above and annotated with prokka [59]. A consensus sequence of the references was generated by mapping the reads of the index isolate (using Snippy). In addition, we filtered variants called for paired isolates by reviewing the alignment of the index isolate reads and excluding positions with a read coverage < 10 and with a fraction of reference allele < 0.5. All variants were manually validated by comparing the alignment of the index isolates and subsequent isolate with the consensus reference. The filtering was performed using SAMtools mpileup, version 1.4 and the alignments were inspected using SAMtools tview, version 1.4 [60].

#### Annotation and functional classification of mutated gene products

The clustering tool CD-HIT, version 4.6.7 [61] was applied to compare proteins whose sequence was altered by mutations confirmed by the approach described above. Unique proteins sequences were annotated by assigning Clusters of Orthologous Groups (COGs) using Reverse Position Specific BLAST (rpsblast), version 2.5.0. The COG database was downloaded from NCBI (ftp://ftp.ncbi.nih.gov/pub/mmdb/cdd/little_endian). Rpsblast results were parsed using the Bioython package [62] Blast, module NCBIXML.

#### Detection of chromosome structural variants

To detect larger deletions, the episode-specific alignment to the complete reference genome was analysed using BEDTools, version 2.26.0 [63] to identify unique intervals within the patient isolates (i.e. not present in all isolates from the same patient) with at least 400 bp read coverage loss. Insertions and smaller deletions were identified by analysis of split reads. Split reads (i.e. reads that can not be represented by a linear alignment and therefore have one or more supplementary alignments as specified in the SAM format available at https://github.com/samtools/hts-specs) were extracted from the episode-specific alignment to the complete reference genome using a python script that is part of the LUMPY framework (https://github.com/arq5x/lumpy-sv). We kept split reads that were unique to one or more isolate per episode and had a breakpoint (defined by start or stop of the read alignment) with a coverage > 10. Breakpoints and read coverage at breakpoint were obtained by parsing the SAMtools mpileup output. Unique split read intervals were confirmed by manual inspection of the alignment, the primary interval and the supplementary intervals were annotated using the complete genome in GFF format and BEDTools. To confirm variants identified with the split reads analysis, we performed a BLAST search for primary and supplementary intervals on the *de novo* assembly graph of the isolates using Bandage, version 0.8.1 [64]. Structural variants were visualised using Geneious, version 8.1.7 (Biomatters).

#### IS256 BLAST search

To obtain the clonal distribution of IS*256*, we performed a blastn search using the IS*256* fasta sequence as a query (downloaded from Isfinder https://www-is.biotoul.fr/index.php) and with the following parameters: minimum coverage 90%, minimum identity 95%, wordsize 32, evalue 0.01. We searched 124 complete genomes available in NCBI repository in May 2017 and 130 draft assemblies of the isolates included in this study.

### Statistical analyses

Statistical analyses were performed in R, version 3.4.1. The chi-square test was used to compare proportions of isolates with at least one mutation and proportion of non-silent mutation among isolate groups. Differences in number of mutations among isolates groups were assessed using the Kruskal-Wallis test. Doubling time and maximum grow rate were calculated by fitting curves using local polynomial regression fitting as performed by the R package *cellGrowth [65].* Enrichment analysis of functional categories among mutated gene products was performed in R by computing the hypergeometric test for each category using the reference genome *S. aureus* TW20 as control.

## DECLARATIONS

### Ethics approval and consent to participate

Human research ethics committee approval was obtained for the study from Austin Health (approval number H2010/04092).

### Availability of data and materials

The datasets supporting the conclusions of this article are available in the European Nucleotide Archive under Bioproject PRJEB22792.

## Authors’ contributions

SGG and BPH designed and planned the study. SGG, SLB, NEH and BPH supplied isolates, clinical data and whole-genome sequencing. SGG, SLB and NEH performed laboratory experiments. SGG, SLB, RG, TS, AGS, MS, RM, NEH, TPS and BPH analysed data. SGG and BPH drafted the manuscript. All authors reviewed and contributed to the final manuscript.

## Acknowledgements

We acknowledge all the patients and health care providers who were involved in the ANZCOSS and VANESSA cohorts. Patients were included in the cohorts by following investigators and centres in Australia and New Zealand: Natasha E. Holmes, Paul D. R. Johnson, Benjamin P. Howden (Austin Health, Heidelberg); Wendy J. Munckhof (Princess Alexandra Hospital, Woolloongabba); J. Owen Robinson (Royal Perth Hospital, Perth); Tony M. Korman (Southern Health, Clayton); Matthew V. N. Sullivan (Westmead Hospital, Westmead); Tara L. Anderson, Sanchia Warren (Royal Hobart Hospital, Hobart); Sally A. Roberts (Auckland District Health Board, Auckland); Sebaastian J. Van Hal (Liverpool Hospital, Liverpool and Royal Prince Alfred Hospital, Sydney); Allen C. Cheng (Alfred Health, Melbourne); Eugene Athan (Barwon Health, Geelong); John D. Turnidge (Australian Commission on Safety and Quality in Healthcare, Sydney). We thank Takehiro Tomita and Susan Ballard (Microbiological Diagnostic Unit Public Health Laboratory, Melbourne) for performing whole-genome sequencing of the ANZCOSS isolates.

## ADDITIONAL FILES

**Additional file 1: Table S1.** Microbiological data and sequence metrics of *S. aureus* isolates included in the study.

**Additional file 2: Table S2.** List of 182 variants identified in 32 isolates (after excluding unrelated strains).

**Additional file 3: Table S3.** List of 81 mutations with predicted modification of protein sequences with COG annotation.

**Additional file 4: Figure S1.** Flow-diagram of the study.

**Additional file 5: Figure S2.** Impact of episode-specific variant filtering (based on read coverage and direct comparison between index and paired isolate alignments) on the total number of mutations identified in paired invasive isolates, when using the closest available complete genome (left panel) and the *de novo* assembly of the index isolate (right panel).

**Additional file 6: Figure S3.** Amino-acid position of variant calls excluded after filtering. Only loci where variants were excluded in at least two episodes are represented.

**Additional file 7: Figure S4.** Number of mutations separating paired invasive isolates to the index blood isolate according to quartiles of the sample collection interval.

**Additional file 8: Figure S5.** Enrichment analysis of COG categories in paired invasive

(A) and colonising isolates (B).

**Additional file 9: Figure S6.** Phenotypic comparison of invasive-colonising pairs. (A) Change in vancomycin MIC from the index isolate to the paired invasive isolate. (B) Waterfall plot of individual changes in vancomycin MIC according to the main treatment before collection of the paired sample. (C) Phenotype convergence in index-paired invasive isolates pairs. (D) Comparison of growth rates (displayed as median, range) within invasive isolates pairs. Pairs with significant increase in doubling time in the paired isolate are indicated with a star. (E) Comparison of cytotoxicity in a subset of paired invasive isolates. Invasive pair from patient 50 exhibited a sharp drop in toxicity and is designated by a star.

**Additional file 10: Figure S7.** Association between number of mutation events and number of IS*256* insertions differentiating paired invasive / colonising isolates from the index isolate among episodes belonging to the ST239 lineage.

**Additional file 11: Figure S8.** Detection of IS*256* in 124 publicly available *S. aureus* complete genomes (panel A) and in the draft assemblies of the 130 isolates included in this study (panel B). For complete genomes the median number of IS*256* copies per sequence type (excluding genomes without IS*256*) is annotated on the right of the bars.

